# A Complete Pedigree-Based Graph Workflow for Rare Candidate Variant Analysis

**DOI:** 10.1101/2021.11.24.469912

**Authors:** Charles Markello, Charles Huang, Alex Rodriguez, Andrew Carroll, Pi-Chuan Chang, Jordan Eizenga, Thomas Markello, David Haussler, Benedict Paten

## Abstract

Methods that use a linear genome reference for genome sequencing data analysis are reference biased. In the field of clinical genetics for rare diseases, a resulting reduction in genotyping accuracy in some regions has likely prevented the resolution of some cases. Pangenome graphs embed population variation into a reference structure. While pangenome graphs have helped to reduce reference mapping bias, further performance improvements are possible. We introduce VG-Pedigree, a pedigree-aware workflow based on the pangenome-mapping tool of Giraffe (Sirén et al. 2021) and the variant-calling tool *DeepTrio* (Kolesnikov et al. 2021) using a specially-trained model for Giraffe-based alignments. We demonstrate mapping and variant calling improvements in both single-nucleotide variants (SNVs) and insertion and deletion (INDEL) variants over those produced by alignments created using BWA-MEM to a linear-reference and Giraffe mapping to a pangenome graph containing data from the 1000 Genomes Project. We have also adapted and upgraded the deleterious-variant (DV) detecting methods and programs of Gu et al. into a streamlined workflow (Gu et al. 2019). We used these workflows in combination to detect small lists of candidate DVs among 15 family quartets and quintets of the Undiagnosed Diseases Program (UDP). All candidate DVs that were previously diagnosed using the mendelian models covered by the previously published Gu et al. methods were recapitulated by these workflows. The results of these experiments indicate a slightly greater absolute count of DVs are detected in the proband population than in their matched unaffected siblings.

## 1 Introduction

Recent advances in genome sequencing technology are improving the accuracy of detecting genetic variants (Wenger et al. 2019). However, the use of a single genome reference for read alignment and variant calling still presents a problem. A sequence mapping algorithm best aligns sequences to a reference when those sequences are present in the reference. Where a sample’s genome deviates significantly enough from the reference, reads will fail to map properly (Sherman et al. 2019). This reference bias can be reduced using pangenome graphs. Pangenome Graphs represent multiple genomes as a series of variants (Garrison et al. 2018). These graphs are further enhanced by incorporating haplotype information that is available in phased genotype datasets. This haplotype information is embedded in a haplotype index (Sirén et al. 2021). In previous work, we have found that mapping error, in both simulation and real-data experiments, is reduced by using population variant data in pangenome graph references (Garrison et al. 2018; Sirén et al. 2021).

Parent-child trios provide evidence of sequence transmission between generations. This helps to identify which variants in the child occurred as de-novo mutations, since these variants will generally be absent in the parents. This information also helps to determine phasing orientation of heterozygotes in the child which can aid in detecting compound-heterozygous candidate DVs. In typical clinical diagnostics, in particular for the case of rare diseases, parental genomes are sequenced to help improve the chances of successful clinical diagnosis of a proband (Clark et al. 2018).

The Undiagnosed Disease Program (UDP) of the National Human Genome Research Institute (NHGRI) is charged with diagnosing previously undiagnosed individuals and discover new variants of clinical significance (Gahl and Tifft 2011; Gahl, Markello, et al. 2012; Gahl, Mulvihill, et al. 2016; Gahl, Wise, et al. 2015; Splinter et al. 2018). In 2009 the UDP started examining cases that have remained undiagnosed after previous exhaustive clinical examination. One part of their process involved sequencing the genomes of patients, including some that included parents, an affected proband and one or more unaffected siblings. Since the beginning of the UDP, they have seen more than 500 different disorders and achieved a diagnostic success rate of over 30 percent, including the discovery of new disorders (Gu et al. 2019). Most of the pediatric cases examined by the UDP over the past 10 years have already had negative diagnostic results from clinical exomes. The UDP applies further technologies, including whole genome sequencing, RNA sequencing, and SNP-chip analysis to more completely explore non-exonic and intergenic regions in an attempt to solve negative exome cases (Gu et al. 2019). One of the more difficult tasks of gene discovery is the detection of variants in highly polymorphic, repetitive, and incompletely-represented regions of the genome, exactly where pangenome graphs can extend accuracy and precision.

In this paper we present VG-Pedigree, a software workflow for mapping and variant calling next generation sequencing data. The workflow leverages pedigrees in genome graphs, and uses machinelearning for variant calling. We also present a companion workflow for identifying candidate deleterious variants which integrates an upgraded implementation of the candidate analysis workflow developed at the UDP (Gu et al. 2019). These upgrades include software portability and usability enhancements, GRCh38 reference compatibility, updates to the population datasets and deleterious predictors used by the previous version of the pipeline, and the incorporation of a new software module that detects mosaicism.

The UDP has applied this workflow to 15 pedigrees that contain a full quartet: one affected child (Proband), one unaffected sibling and two unaffected parents. Of the probands that had a known diagnosis, the corresponding DVs that were detected based on the mendelian models covered by the previously published candidate analysis workflow and were recapitulated by this workflow. The workflows are modular such that the candidate analysis workflow can be run by any set of alignments in the BAM format and a joint-called pedigree dataset in the VCF format.

## 2 Results

### 2.1 Overview of VG-Pedigree

VG-Pedigree goes through a number of stages before final variant calling (Fig. 1A). First, the set of short reads in the parent-parent-child trio of the pedigree are mapped to a pangenome graph reference based on the 1000 Genomes Project dataset, termed 1000GP, using VG Giraffe, and variants are then called using *DeepTrio* (Fig. 1B) (Auton et al. 2015; Sirén et al. 2021). Next, variants in 1000 Genomes Project haplotypes that appear missing in the DeepTrio-called variants are imputed. The resulting variant file is phased using both alignment and pedigree information (Fig. 1C). A parental graph reference is then constructed using only the parental genotypes from the joint-called VCF file (Fig. 1D). A haplotype index of this graph reference for VG Giraffe is generated from the phased genotypes of the parental samples. Once this graph is constructed, the proband and siblings reads are re-mapped to this new parental graph reference and variants are re-called using the new mappings (Fig. 1B). Finally, the newly-called variants of the child and sibling samples are joint-called with the old parental variants to form the final joint-called pedigree VCF.

**Figure 1.**
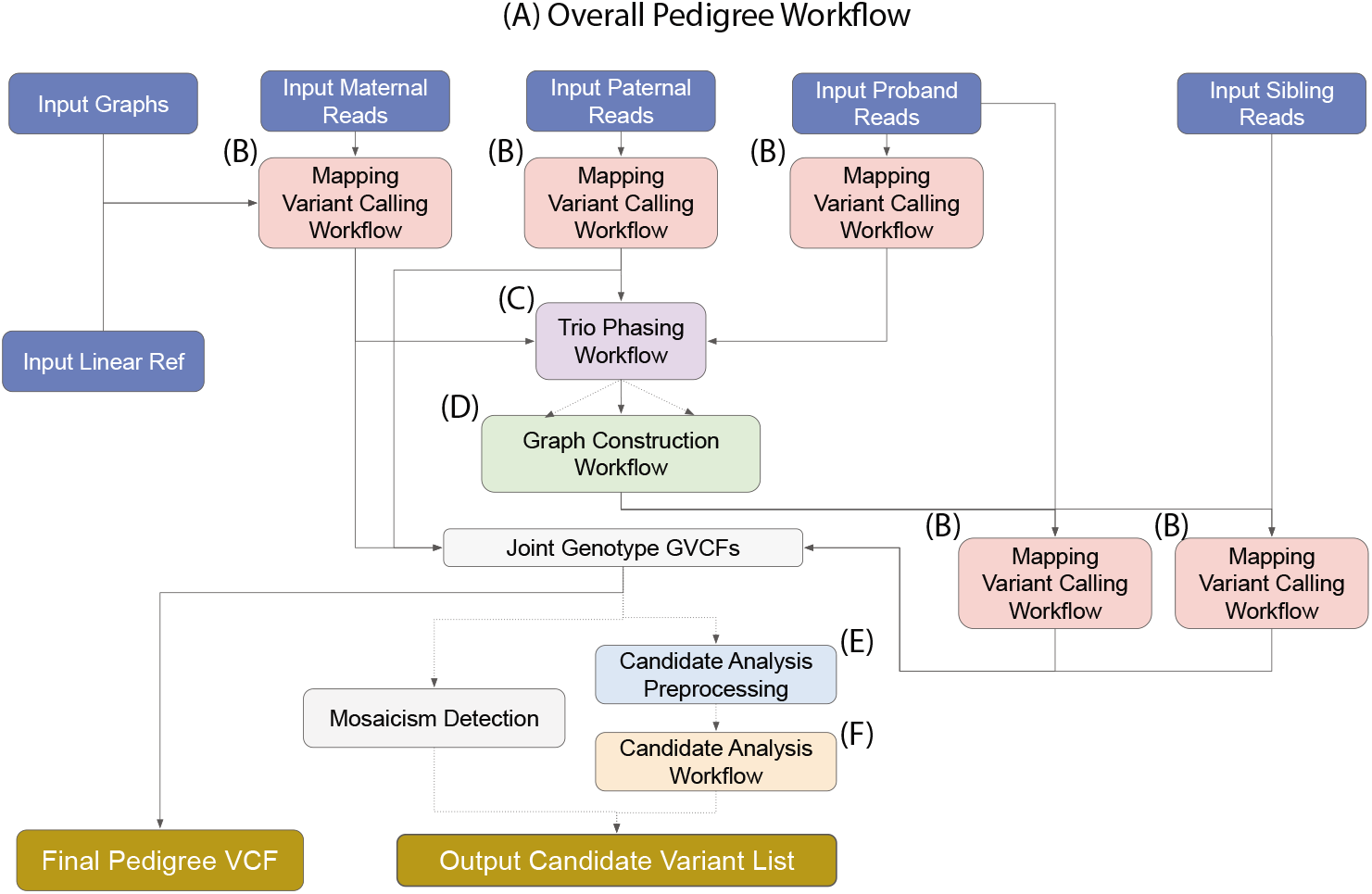
Toil-VG Pedigree workflow. Dotted lines indicate optional pathways in the workflow. (A) Overall workflow diagram. (B) Single sample alignment and variant-calling workflow. (C) Trio joint-genotyping and phasing workflow. (D) Parental graph construction workflow. (E) Workflow for preprocessing and annotation of pedigree variants required for candidate analysis. (F) The candidate analysis workflow.

The candidate analysis workflow takes as input the set of alignments and variant calls from VG-Pedigree and outputs a final set of candidate DVs for the proband. This is done through a series of filters and annotations. First, SnpEff is used to annotate the type and function of variants within the joint-called pedigree VCF file (Cingolani et al. 2012). The deleteriousness of these variants is predicted using the Combined Annotation Dependent Depletion (CADD) software tool (Rentzsch et al. 2021) (Fig. 1E). Next, a series of filtration and analysis methods are applied to the annotated variants and the workflow outputs a set of candidate DVs for the proband (Fig. 1F). The methods applied in the candidate analysis workflow are an implementation of the methods described in the Gu et al. study (Gu et al. 2019). In this paper, we present enhancements to the methods and software of the candidate analysis workflow. An additional module of the analysis workflow has also been developed which automatically detects the presence and type of mosaicism in the designated proband. These methods and improvements together provide a more complete and accurate dataset from which to discover rare variants that are causal to genetic diseases over the previous iteration.

We evaluated performance of this workflow based on four main metrics. First, we evaluated the ability of the workflow to accurately align reads to the correct position in a genome. Second, we assessed the accuracy of variant calls based on those alignments. Thirdly, we looked at the ability of the analysis workflow to capture DVs in the proband population versus the unaffected sibling population. Finally, we examined the runtime and costs of running this workflow using a commercial cloud environment.

### 2.2 Mapping Evaluation

Mapping was evaluated with both simulated and real sequencing data. The former considers measures of mapping reads with a known position. This was done by simulating reads from haplotypes whose corresponding path locations in the graph are known, so that we could identify when a read was mapped to the correct location on the graph (see supplementary methods S1). Figure 2 illustrates the performance of 10 million read pairs that are simulated from the Genome-in-a-Bottle (GIAB) HG002 version 4.2.1 high confidence variant sets (Zook 2020). We also examined stratified performance across regions of interest using 100 million reads simulated from the GIAB high confidence regions. These regions were all defined by GIAB: (the Global Alliance for Genomics and Health Benchmarking Team et al. 2019) low complexity regions that comprise regions of low sequence variability, low mappability regions that are made up of duplicated and paralogous sequence, the Major Histocompatibility Complex (MHC) which is known for maintaining a high density of variation, 1000GP variant regions excluded from the GIAB sample (1000GP-excluded), and, specifically for HG002, the complex medically relevant genes (CMRG) included in a study by Wagner et al. (Wagner et al. 2021).

**Figure 2.**
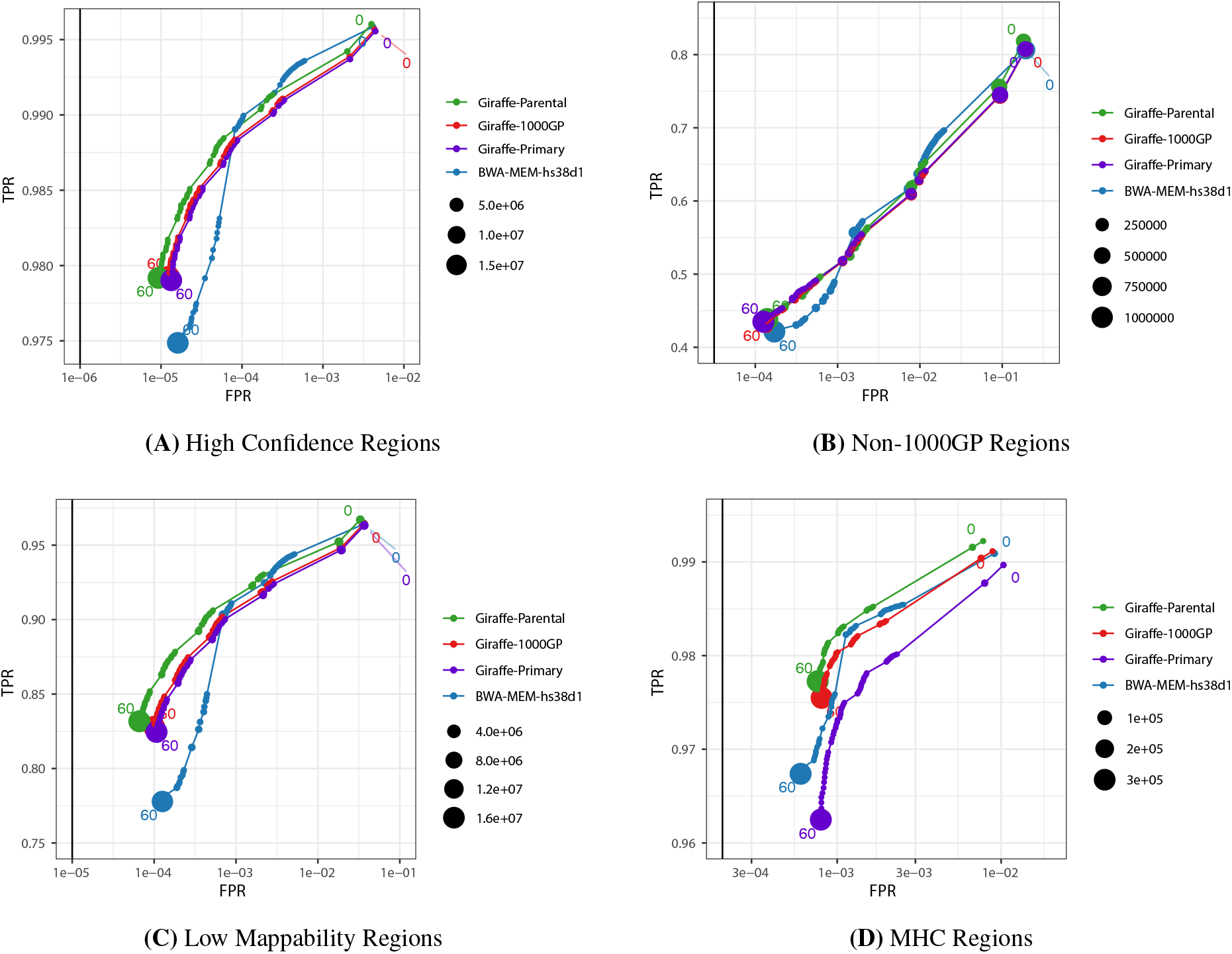
Mapping performance of 100 million read pairs simulated from HG002 high confident datasets. Four different alignments are compared across four different regions and ROC curves are plotted with a log-scaled false positive rate on the x-axis and a linear-scaled true positive rate on the y-axis with the mapping quality as the discriminating factor. Green curves represent graph alignments against the parental graph reference constructed from HG003 and HG004 illumina read graph alignments. Red curves represent alignments against the 1000GP graph reference. Purple curves represent alignments to the primary GRCh38 linear graph reference. Blue curves represent linear alignments against the `hs38d1` reference using bwa-mem. (A) Alignments in GIAB v4.2.1 confident regions (from 1 million simulated read set). (B) Alignments in non-1000GP confident regions (from 1 million simulated read set). (C) Alignments in GIAB v4.2.1 low mappability regions (from 100 million simulated read set). (D) Alignments in GIAB v4.2.1 MHC regions (from 100 million simulated read set).

All conditions evaluated consist of the combination of a mapper and a reference (see supplementary methods S2). The *Giraffe-Parent* condition used VG Giraffe (Sirén et al. 2021) to align reads to the parental graph reference as produced by the workflow up to graph construction (Fig. 1D). The *Giraffe-1000GP* condition used VG Giraffe to align reads to the pangenome reference. The *Giraffe-Primary* condition used VG Giraffe to align reads to a linear graph reference as produced using only the `hs38d1` reference with no variation. And the *BWA-MEM-hs38d1* condition used BWA-MEM (H Li 2013) to align reads to the `hs38d1` human reference genome.

Figure 2 shows the receiver-operator-curves (ROC) of each tested mapper in all confident regions, 1000GP-excluded regions, low mappability regions, and MHC regions. The curves are stratified by mapping quality (MAPQ). In each evaluated region, *Giraffe-Parent* produced the highest F1, both for reads with MAPQ60 and across all reads. When looking at 1000GP-excluded variants within stratified regions, *Giraffe-Parent* produced the highest total F1 across low complexity regions (Fig. S1), low mappability regions (Fig. S2), MHC regions (Fig. S3), and CMRG regions (Fig. S4).

For all GIAB high confidence regions, *Giraffe-Parent* gave the most accurate alignments relative to the other examined mappers. *Giraffe-Parent* also achieved the highest total of correctly mapped reads in all but the CMRG regions, the highest total of reads mapped at MAPQ60 in low mappability MHC and CMRG regions, and the highest average percent identity between aligned reads and the reference sequence across all regions (Table S1). In the high confident 1000GP-excluded regions of the HG002 sample, *Giraffe-Parent* achieved the highest proportion of correctly mapped reads, MAPQ60 reads, and average sequence identity (Table S2). *Giraffe-Parent* also produced the highest proportions of perfectly-aligned and gaplessly-aligned reads, and the lowest proportion of soft-clipping reads across all examined confident (Table S3) and 1000GP-excluded regions (Table S4).

### 2.3 Variant Calling Evaluation

In addition to examining the mapping performance of the workflow we measured the accuracy of variants called in each workflow. Here we use the version 4.2.1 release of the HG001, HG002 and HG005 truthset benchmarks as published by GIAB (Zook 2020). RealTimeGenomics `vcfeval` tool (Cleary et al. 2015) and Illumina’s `hap.py` haplotype aware variant comparison tool (*Illumina/hap.py* 2020) were used when comparing the results of variants called using alignments of real reads to various combinations of mappers and references. The mappers and references used include VG Giraffe against the 1000GP graph (termed *Giraffe-1000GP*) VG Giraffe against the parental graph (*Giraffe-Parent*), `BWA-MEM` against the linear `hs38d1` reference (*BWA-MEM-hs38d1*), and Illumina’s `Dragen` platform version 3.7.5 (Illumina 2020; *Hidden Treasures - Warm Up precisionFDA* 2020; *Truth Challenge V2: Calling Variants from Short and Long Reads in Difficult-to-Map Regions precisionFDA* 2020) against the linear `hs38d1` reference (*Dragen-hs38d1*) (see supplementary methods S3 and S4).

We used *DeepTrio* version 1.1.0 with trained child and parent models for variant-calling comparison in HG002 and HG005 samples. Training used the Ashkenazi (HG002, HG003, HG004), Han Chinese (HG005, HG006, HG007) and CEPH (HG001, NA12891, NA12892) trio alignments using the *Giraffe-1000GP* method for model training (see supplementary methods S4.2). We left out chromosome 20 for validation purposes. Supplementary Figure S10C-D and Supplementary Table S5 show the results of training for HG002 and Supplementary Figure S11C-D and Supplementary Table S6 for HG005 results. The total number of errors in chromosome 20 reduced from 1,070 to 1,051 (1.78%) and from 1,130 to 909 (19.56%) variants for HG002 and HG005, respectively.

We then assessed HG001 using the same training method for the model used in evaluating HG002 and HG005. The models were re-trained with Girafe-1000GP-aligned read data while the HG001, NA12891 and NA12892 samples were completely held out for validation purposes. The DeepTrio-called variants achieve the highest accuracy (F1: 0.9976) using *Giraffe-Parent* (Table 1A-B). This represents a total variant error (false positive and false negative) reduction of 4,844 variants between *Giraffe-Parent* and *BWA-MEM-hs38d1* relative to an error reduction of 2,925 variants between *Giraffe-1000GP* and *BWA-MEM-hs38d1*. In the 1000GP-excluded variants, the *Giraffe-Parent* accuracy (F1: 0.9748) outperforms *Giraffe-1000GP* (F1: 0.9717) by a greater margin than *Giraffe-1000GP* outperforms *BWA-MEM-hs38d1* (F1: 0.9691). This reflects an error reduction of 3,210 variants between *Giraffe-Parent* and *BWA-MEM-hs38d1* relative to an error reduction of 1,481 variants between *Giraffe-1000GP* and *BWA-MEM-hs38d1*.

**Table 1.**
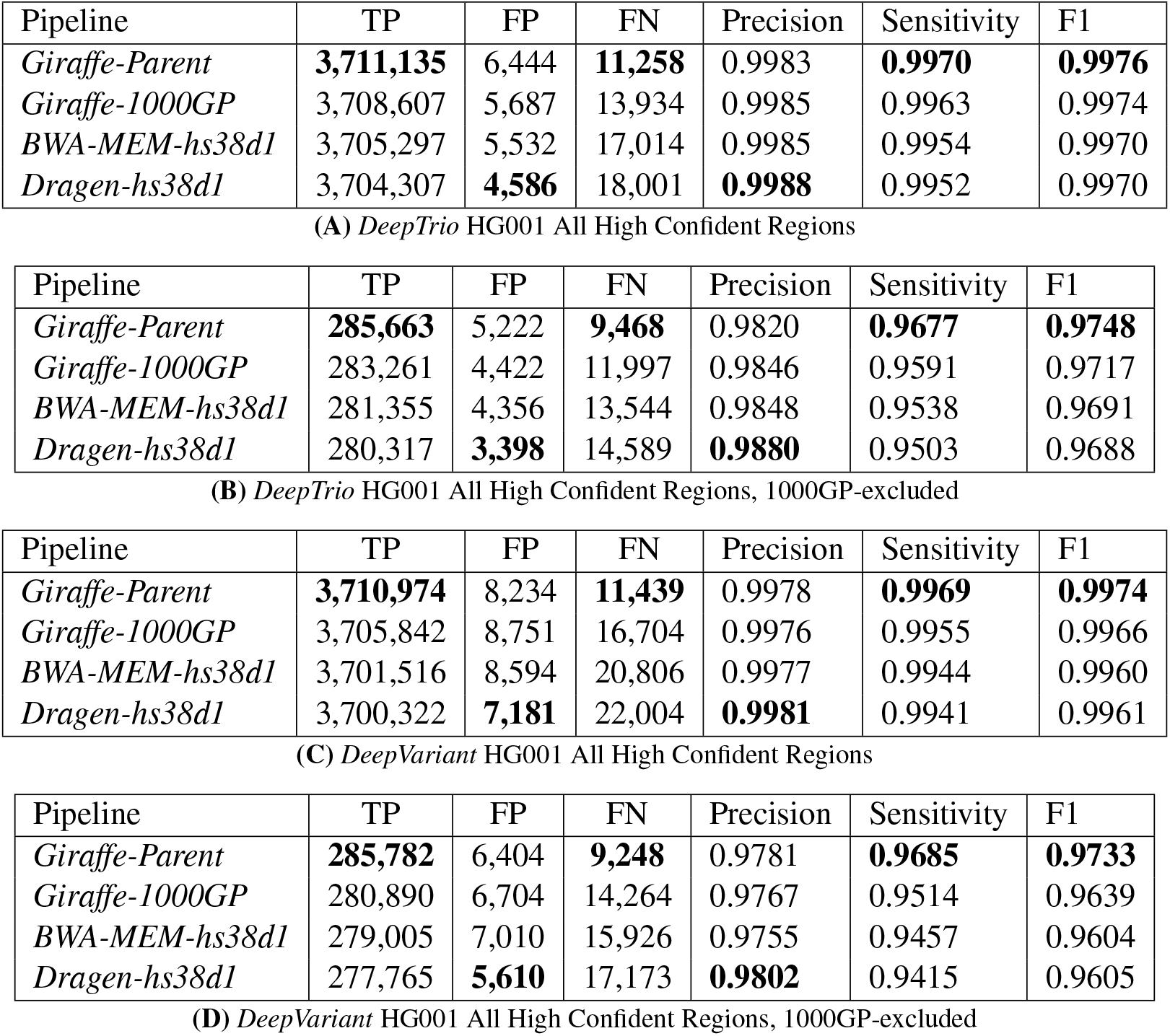
VCFeval HG001 *DeepTrio* and *DeepVariant* Performance. VCFeval performance of the graph-based and linear-based pipelines with respect to HG001 GIAB v4.2.1 truth variant call sets stratified by (A) *DeepTrio* on all HG001 regions, (B) *DeepTrio* on HG001 regions excluding 1000GP variants, (C) *DeepVariant* on all HG001 regions, and (D) *DeepVariant* on HG001 regions excluding 1000GP variants. All mapped reads were called using *DeepTrio* and *DeepVariant* v1.1.0 genotyper using trained models. Best values in each column are highlighted in bold text.

We also tested *Giraffe-Parent* using the default *Deep-Trio* version 1.1.0 models, which were not trained with Giraffe alignments. We found that in using the HG005 and HG002 trios *Giraffe-Parent* or *Giraffe-1000GP* with the default *DeepTrio* models outperforms the results achieved using standard BWA-MEM (Table S7A-B). The same performance gains are observed for *Giraffe-Parent* in more difficult regions for both HG002 and HG005 samples (Tables S8 and S9).

ROC curves for DeepTrio calls stratified by genotype quality also show performance gains. Figure 3 shows the ROC curves between the graph-based and linear-based alignment methods in HG001 for all confident regions and 1000GP-excluded variants respectively. Supplementary Figures S10A-B and S11A-B illustrates performance in the same regions but for the HG002, and HG005 samples using the default DeepTrio models, respectively.

**Figure 3.**
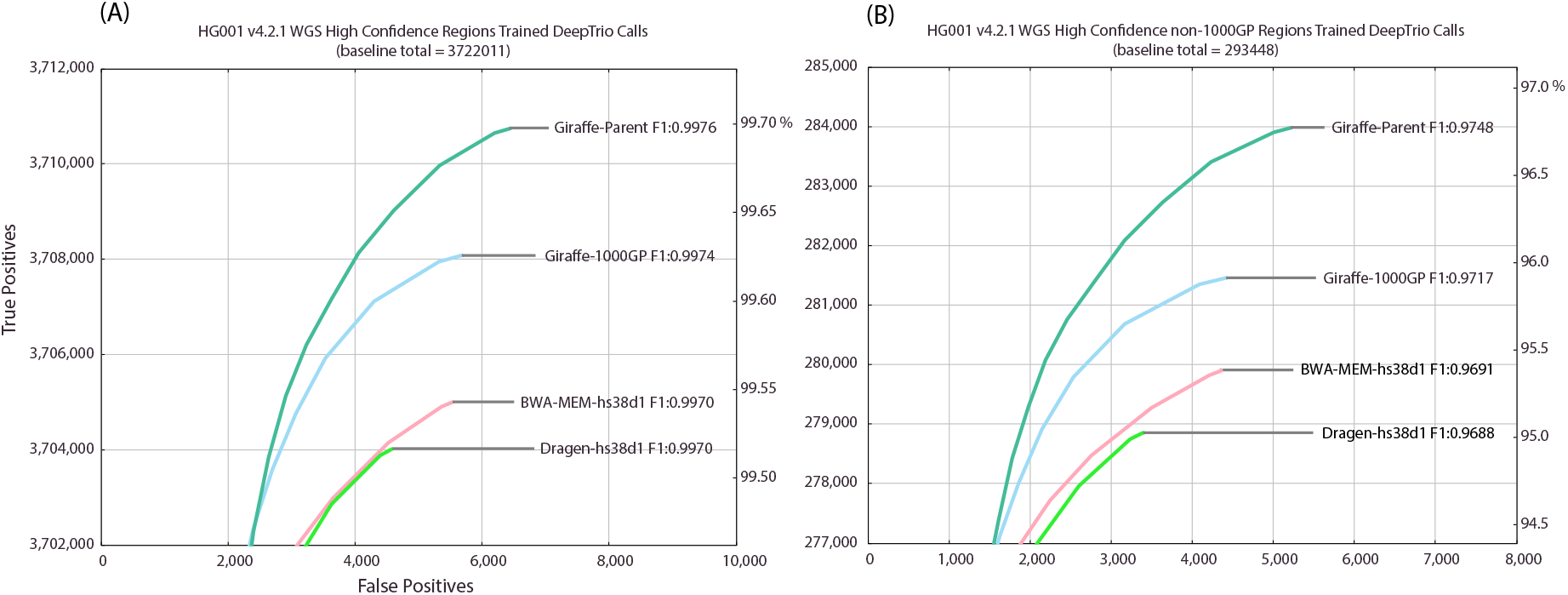
ROC curves of *DeepTrio* variant calling performance of the graph-based and linear-based pipelines with respect to HG001 GIAB v4.2.1 truth variant call sets stratified by (A) HG001 high confident whole genome regions using trained *DeepTrio* models, (B) HG001 high confident whole genome regions excluding 1000GP variants using trained *DeepTrio* models.

To compare the results of our trio-based pipeline to other calling methods we additionally tested the non-trio based *DeepVariant* version 1.1.0 and Illumina’s *Dragen* platform version 3.7.5 variant callers, using Giraffe pangenome and BWA-MEM linear reference mappings. This evaluation assesses gains in variant calling accuracy brought by using trios vs. simply mapping to a linear reference, a population based pangenome graph (*Giraffe-1000GP*), or to a graph containing the subject’s parental information (*Giraffe-Parent*). Dragen, which uses an algorithm similar to that of GATK HaplotypeCaller, was used to call variants against the `hs38d1` reference for each set of alignments. Once again, *Giraffe-Parent* produced the most accurate variant calls for HG002 and HG005. *Giraffe-Parent* produced the highest F1 score (0.9965) in all confident regions for HG002 (Table S10). This is in contrast with the F1 performance of *Giraffe-1000GP* (0.9953) and *BWA-MEM-hs38d1* (0.9940). Total error is reduced by 18,754 variants between *Giraffe-Parent* and *BWA-MEM-hs38d1* relative to an error reduction of 9,995 between *Giraffe-1000GP* and *BWA-MEM-hs38d1*. For HG005, *Giraffe-Parent* produced the highest F1 score (0.9958) in all confident regions (Table S11). This is in contrast with *Giraffe-1000GP* (F1: 0.9944) and *BWA-MEM-hs38d1* (F1: 0.9931). Total error is reduced by 20,724 variants between *Giraffe-Parent* and *BWA-MEM-hs38d1* relative to an error reduction of 10,489 between *Giraffe-1000GP* and *BWA-MEM-hs38d1*.

During evaluation of DeepVariant calls, like in our DeepTrio evaluations, we focused on using models that were not trained with pangenome graph alignments of the samples used in evaluation. For HG001 alignments, a trained DeepVariant model was used in evaluating HG001 whole genome results. This model was trained using just the *Giraffe-1000GP*-aligned HG002 and HG004 sample reads. For evaluations of DeepVariant calls on HG002 and HG005 alignments, the default models of DeepVariant version 1.1.0 were used. In HG001, the *Giraffe-Parent* method achieves the highest accuracy (F1: 0.9974) representing a total variant error reduction of 9,727 variants between *Giraffe-Parent* and *BWA-MEM-hs38d1* relative to an error reduction of 3,945 variants between *Giraffe-1000GP* and *BWA-MEM-hs38d1* (Table 1C). For HG002, the DeepVariant-called variants also achieves the highest accuracy (F1: 0.9970) using *Giraffe-Parent* (Table S7C). This represents a total variant error reduction of 4,849 variants between *Giraffe-Parent* and *BWA-MEM-hs38d1* relative to an error reduction of 131 variants between *Giraffe-1000GP* and *BWA-MEM-hs38d1*. Finally, for HG005, the *Giraffe-Parent* method produces the highest accuracy (F1: 0.9966) representing a total variant error reduction of 8,009 variants between *Giraffe-Parent* and *BWA-MEM-hs38d1* relative to an error reduction of 2,063 variants between *Giraffe-1000GP* and *BWA-MEM-hs38d1*. (Table S7F).

Breaking down the analysis to SNPs and INDELs reveals the same trend. The *Giraffe-Parent* produced the highest F1 scores in HG002 in all examined regions except for the CMRG genes, where *Dragen-hs38d1* achieves a higher accuracy in INDELs (F1: 0.959108) relative to *Giraffe-Parent* (F1: 0.958785) (Tables S12 S13 S14 S15 S16, and S17). Similar statistics are observed in HG005, where *Giraffe-Parent* alignments produce the highest F1 in all SNPs and INDELs across all confident regions (Tables S18 S19 S20 S21 S22).

### 2.4 Candidate Analysis Evaluation

The last evaluation investigates the workflows ability to identify DVs that are relevant to clinical disorders. We ran the workflow on 15 nuclear pedigrees of at least 4 individuals in size. Out of the UDP set of 50 cohorts with identified candidate variants, a set of 15 cohorts were randomly chosen. The 15 cohorts include 15 probands and 22 unaffected siblings comprising 18 females and 19 males. 11 of the UDP probands from these cohorts that have a known genetic diagnosis had their causal variants recapitulated by this workflow. The list of Mendelian models detected include homozygous recessive, de-novo, hemizygous, X-linked, mitochondrial, and compound-heterozygous genotypes. Of the 12 examined probands that have a diagnosis attributed to a CLIA-validated variant, 5 were identified with de-novo dominant non-synonymous changes in an exonic region, 2 had a de-novo dominant frameshift in an exonic region, 2 had compound-heterozygous variants where both were non-synonymous changes in exonic regions, and 1 had a compound-heterozygous variant with a non-synonymous change in an exon and a change in an intronic/splice-site region. Supplementary Table S23 shows the number and type of candidate variants detected by the workflow for all 37 individuals.

We compared the number and type of clinically-relevant variants that are identified between the affected proband population and their matched unaffected sibling population to indirectly evaluate the pipelines ability to identify DVs. This analysis runs in two steps. First, for each family, the affected offspring are set as the proband in the workflow and the unaffected offspring are set as the unaffected siblings. Then for the second step, for each family, the unaffected offspring are set as the proband and the affected offspring are set as the unaffected siblings. Finally, the set of candidate DVs from the probands in the first step are compared against the set of candidate DVs from their matched unaffected siblings in the second step.

There is an expected baseline load of rare deleterious variants that all individuals inherit due to de-novo mutation and inefficient selection against segregating variants (Henn et al. 2015). Given a large enough sample set, we expect the median number of rare deleterious variants in the proband population to be slightly different from the median number of rare deleterious variants in the unaffected sibling population. Due to two factors, the probands level of genetic burden is hypothesised to be slightly greater than that of their unaffected siblings: all probands in this analysis currently show phenotypic expression of their disease, and the unaffected siblings are of similar age.

Figure 4A shows the distribution between these two populations in the 15 pedigree cohort sample set. Figure 4B shows the distribution of differences in the candidate DVs between the matched proband and siblings. X-linked recessive candidate DVs were excluded from both populations in order to improve comparability between male and female samples. Compound-heterozygous candidate pairs and candidate alleles that occupy the same locus are also counted as one candidate for the purposes of this comparison. The number of candidate DVs in the proband population are significantly different from their matched unaffected siblings set of candidate DVs (Wilcoxon signed-rank test p-value=0.03).

**Figure 4.**
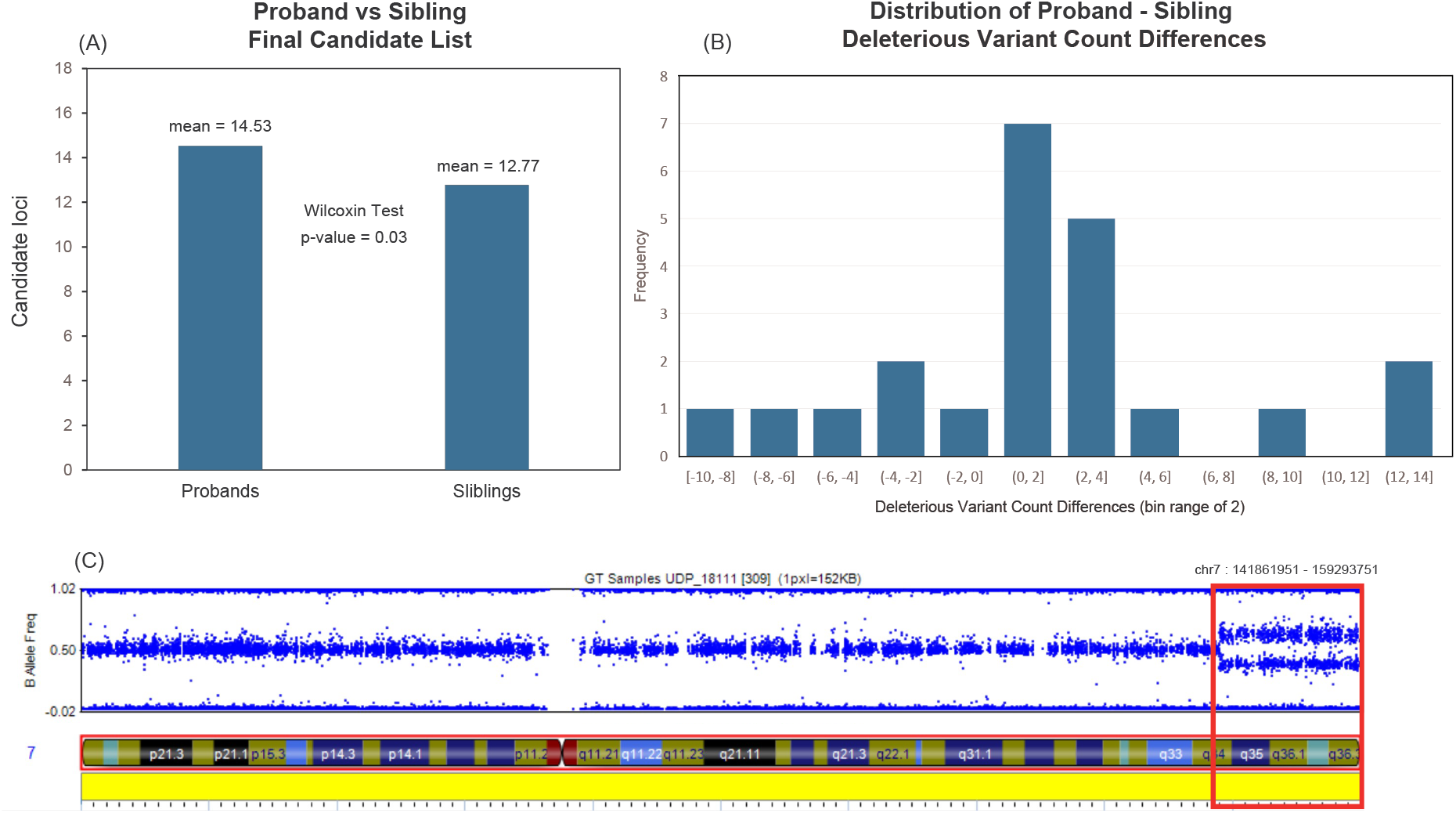
Proband-sibling pairwise candidate analysis results on 15 nuclear families of at least quartet in size, comprising a population of 15 probands and 22 siblings. Plot (A) shows the average number of candidate variants between the probands and sibling populations. The proband population holds an average of 14.53 CVs while the sibling population has an average of 12.77 CVs. A one-tailed Wilcoxon signed-rank test of the hypothesis that the probands have greater numbers of candidate variants than their matched siblings produced a p-value of 0.0333. (B) The distribution of proband - sibling candidate variant list size differences. (C) A mosaic region identified by the workflow (red box) overlaid with the snp-chip B allele frequency plot for a UDP sample.

In addition, we ran the workflow on 4 undiagnosed cases that have previously shown a negative or inconclusive clinical exome and negative commercial genome assay results. From these samples, we have produced a number of candidate DVs. Of the 4 cases, 2 have candidate DVs that match their phenotypic profile and are being examined for clinical function, the other 2 cases are undergoing further investigation. One of the 2 cases had an identified mosaic region on chromosome 7 detected by the candidate analysis workflow (Figure 4C). Further phenotypic and candidate DV details are omitted to protect patient privacy. Supplementary Tables S24 and S25 shows the number and type of candidate variants detected by the workflow.

### 2.5 Runtime Evaluation

The workflows examined are runnable on the Terra platform (*Terra* 2021). When running on a quartet with 30 to 35x coverage paired read data, the workflow takes a little more than 8000 CPU hours for a total cost of approximately $100 (Table S26). The VG-Pedigree pipeline makes up the majority of the computation at about 8000 CPU hours and costs $92-95, while the candidate analysis workflow runs in about 200 CPU hours costs $3-5. Costs can vary based on the load of the cloud compute system and the availability of lower-cost preemptable nodes.

## 3 Discussion

There is growing evidence that rare variants have the effect sizes, diversity and abundance necessary to explain a substantial portion of human genetic load (Hernandez et al. 2019; Simons et al. 2014; X Li et al. 2017). Pedigrees can help resolve harder-to-study regions by giving orthogonal evidence in the form of Mendelian inheritance to enhance the statistical power and phasing accuracy to categorize compound-heterozygote and de-novo variation from a list of called variants (Shugart et al. 2012; Peng et al. 2013; Roach et al. 2010; Sul et al. 2016). Graph-based approaches leverage additional variation information during read mapping to mitigate the problems of alignment to complex regions of the genome (Garrison et al. 2018; Sirén et al. 2021). The methods and software developed in this project are designed to maximize the biological information available to detect and interpret individual-level variation. The software developed is scalable so that it can easily run on high performance compute clusters that support common batch systems like Slurm (Yoo et al. 2003) or Kubernetes (*Production-Grade Container Orchestration* 2021). It is publicly accessible in the `toil-vg` GitHub repository and in `WDL` format which is published in the Dockstore repository (O’Connor et al. 2017; Voss et al. 2017).

Alignment performance of short-sequenced reads is improved across all examined confident regions in the GIAB samples. This translates to better coverage, mapping quality and greater variant calling accuracy in these regions. All examined UDP cases that have a known genetic diagnosis based on the Mendelian models covered by the previously published candidate analysis workflow (Gu et al. 2019) have their causal variants recapitulated by this workflow. The candidate analysis evaluation indicates detectable differences in the number of candidate DVs identified between the affected and unaffected offspring populations. This result shows a similar trend to that of the analysis done on exome datasets from a larger sample set, which also showed a statistically significant difference (Gu et al. 2019). The main improvement in this analysis over the previous analysis is that this analysis covers the whole genome including intronic and intergenic regions.

A number of areas can be improved within this workflow. One example is the training model used in DeepTrio. Our training used a very limited number of benchmark samples, which was limited further to leave benchmark data for testing and development. Given these limitations, there is room to improve the *DeepTrio* model when additional well-sequenced and diverse benchmark samples become available.

Variant calls from graph-based alignments are prone to error due to the conversion of the native graph alignment map (GAM) format output from VG alignments to the linear reference BAM format. Information about the exact path of reads is lost during this projection step which can result in reads appearing different from the linear reference genome when the variant is already present in a path in the graph reference.

Structural Variants are an important component to the set of rare variants that contribute to disease (Weischenfeldt et al. 2013; Abel et al. 2020). In previous work, there have been efforts to tailor pangenome graphs and variant caller algorithms to improve the accuracy of detecting structural variants (Sirén et al. 2021). Another avenue to improve this workflow is to apply pangenome graphs with incorporated structural variant information as a module that runs concurrently to the VG-Pedigree workflow.

Further runtime improvements could also be made. The workflow takes about 1.5 days and approximately 8000 total CPU hours at a cost of about $100 to process one family. This is moderately expensive and slow relative to traditional methods, which have well-tuned hardware acceleration solutions and years of work optimizing computation time. GPU acceleration or field-programmable-gate-array (FPGA) implementations of the graph alignment algorithm could substantially accelerate the computation of the graph-based algorithms.

There are a number of refinements that could be made to the most expensive parts of this workflow. Reference construction of the parental graph could be improved by altering and pruning the haplotype index with the haplotypes discovered by the trio-backed phasing stage of the pipeline. The use of graphbased variant callers would remove the need to surject alignments to linear BAM files and therefore maintain potentially more information that could be used to produce more accurate calls.

New pangenome graphs are continuously being updated and tested as more population variation is characterized. The Telomere-to-Telomere genome project (T2T) has recently released a genome reference which exhaustively captures the centromeric and telomeric sequence better than the previous GRCh38 version of the human genome (Nurk et al. 2021). The Human Pangenome Reference Consortium (HPRC) is a group of research institutions that are tasked with the development of a pangenome reference using the latest methods and data. By characterizing regions of the genome not well represented by existing variant datasets, the pangenome references developed by the HPRC that incorporate new T2T sequences should further improve the performance and accuracy of the workflows presented in this paper.

## 4 Methods

### 4.1 VG Pedigree Workflow

Pangenome graphs provide a framework for leveraging genomic variation information to create a better-informed mapping procedure than that provided by a linear genomic reference. The workflow presented here goes through a number of stages (Fig. 1A). The first stage establishes parental haplotypes to construct a parental-backed graph reference. It takes short reads from a trio and aligns each to a population-informed graph reference. We use a graph based on the 1000 Genomes dataset (Auton et al. 2015; Sirén et al. 2021). It is still the largest and most diverse set of phased genotypes available to the public with broad consent. The 1000GP graph is based on the `hs38d1` human reference genome and the 1000 Genomes Project phase 3 variant set, and is available in a publicly accessible Google Cloud bucket.

Alignment of the parent-child trio to the 1000GP graph goes through a number of steps that split and merge read alignments to enable distributed computation (Fig. 1B, Supplementary Fig. S5). This greatly reduces time spent aligning reads, which is a major bottleneck for the workflow. Afterwards, each chunked alignment is projected back to the linear genome reference coordinate space and corrected for duplicates and missing mate information and indels are realigned using ABRA2 (Mose et al. 2019). Following alignment, samples in the trio are variant-called, producing a per-sample gVCF genotype called file. A trio-based *DeepVariant* extension (Poplin et al. 2018), Google’s *DeepTrio* (Kolesnikov et al. 2021), is used to call variants in this workflow. *DeepTrio* first generates images based on the alignments between the parent and child reads. Then the *DeepTrio* variant caller is run concurrently to call gVCFs for each contig for each sample in the trio. The gVCFs are next joint-called with the Glnexus package (Yun et al. 2020) in order to merge and recall potentially uncalled variants in the trio. Joint-calling gVCFs enhances DeepVariant-based calls by reexamining trio variant sites that were confidently called in one sample but not another. The joint-called trio VCF is then divided by autosomal and sex-chromosomal contigs, with the mitochondrial contig only preserving the maternal set of called genotypes and the Y chromosomal contig preserving the paternal set of called genotypes.

A number of different schemes for phasing these variants were explored using combinations of *Eagle* (Loh et al. 2016), *WhatsHap* (Martin et al. 2016), and *SHAPEIT4* (Delaneau et al. 2019). Supplementary Table S27 illustrates the performance of combinations of these programs when phasing the GIAB HG002 sample. Supplementary Table S28 shows phasing performance for the GIAB Ashkenazi trio with respect to GRCh38- or GRCh37-based graph alignments. Using *Eagle* followed by *WhatsHap* produced the largest blocks of phased variants while maintaining a switch error rate close to, or better than, the method with the largest median haplotype block size from this list: *WhatsHap* in combination with *SHAPEIT4*. Following the alignment and variant calling step, a phasing sub-pipeline is run on these contig VCFs using the Eagle-WhatsHap phasing method (Fig. 1C, Fig. S6). Missing genotypes are imputed using *Eagle* version 2.4.1 (Loh et al. 2016). Finally the contig VCFs are phased with trio- and read-backed methods using *WhatsHap* (Martin et al. 2016). That final set of contig VCFs are then filtered down to just the parental genotype sets and passed into the graph construction workflow.

Following the phasing stage of the workflow, the phased variants from that step and a linear reference in FASTA are passed as input into the graph construction step (Fig. 1D, Fig. S7). VG mappers use a variety of indexes (Sirén et al. 2021). To facilitate this need, the construction workflow generates a combination of indexes based on the requirements of the VG Giraffe mapper.

After constructing the parental graph, the offspring reads can be realigned to it. GVCFs are called from offspring alignments to the parental graph reference. (Fig. 1B, Fig. S5). Finally, variants are jointly called, once again with the Glnexus package (Yun et al. 2020), by combining previously-computed gVCFs of the 1000GP-aligned parents with gVCFs derived from the parental graph-aligned offspring.

The methods developed here for the VG-Pedigree workflow are implemented in the software framework toil-vg under the ‘toil-vg pedigree‘ subcommand which makes use of the TOIL workflow engine (Vivian et al. 2017) for cloud-based and cluster-compute systems and is available on GitHub at https://github.com/vgteam/toil-vg. The workflow is also made available in `WDL` format in the Dockstore (O’Connor et al. 2017; Voss et al. 2017) repository at https://dockstore.org/workflows/github.com/vgteam/vg_wdl/vg-pedigree-giraffe-deeptrio:master.

### 4.2 Candidate Analysis Workflow

A primary endpoint goal for this workflow is variant detection to identify likely causes of the genetic disorders in the UDP cases. Traditional variant filtration techniques narrow down a set of variants, but they are usually not exhaustive enough to narrow the list down to an actionable number of variants without truncation (Pedersen et al. 2021; Kobren et al. 2021). Further, they often do not specialise in the detection of compound-heterozygous candidates in non-coding regions. Traditionally, a large proportion of work is needed to validate the clinical functionality for each variant (Baldridge et al. 2017). Given this downstream cost, this workflow focuses on reducing that cost by minimizing the number of variants that need to be examined in the final list. The analysis workflow takes in a very large set of variants and filters them by examining a series of variant attributes each of which follows an order of most-certain to least-certain true-positive data types (Fig. 1F).

Additional improvements and features were added to this implementation of the methods developed in the Gu et al. study. In this paper, we have adapted all components and annotations used by the workflow to be compatible with the GRCh38 reference genome coordinate system. The CADD engine software suite has been updated to version 1.6 which incorporates greater accuracy in determining deleterious variants located in splice sites and introns (Rentzsch et al. 2021). We have also updated the population annotation dataset to use GnomAD v3.1 which has incorporated a larger proportion of samples producing more accurate and exhaustive population allele frequencies (Karczewski et al. 2020). The maximum minor allele frequency (MAXMAF) calculation implemented in the `population/deleterious-backed variant filtration` module was altered to use a binomial instead of a poisson distribution (Fig. S8D). A critical bug was patched that was found to erroneously output X-linked candidate variants for females. We implemented a new module that automatically detects the presence, location and type of copy number variant (CNV) mosaicism in the proband.

The alignment and variant calling workflow output is processed with various annotation programs before they are able to be passed as input into the candidate analysis workflow. Post processing the final datasets comprises SnpEff annotation, indel-realignment, and converting to a one variant per row format that has pedigree consistent indels, for each of the samples in the pedigree (Fig. 1E, Fig. S9). The CADD (Rentzsch et al. 2021) software suite is used in this analysis workflow to predict the deleteriousness of a given variant. Any variants that are unique to the CADD database in the joint VCF have a deleterious score calculated by the software.

The analysis portion of the workflow examines and filters the pedigree variant file in the context of Mendelian inheritance, alignments against the parental-based graph reference, population variant frequency, and predictions of variant effects on gene function and expression (Gu et al. 2019) (see supplementary methods S5). Using these filters generates a set of variants that are further filtered by examining the BAM files for sequence and alignment noise surrounding each variant (Gu et al. 2019). This produced a final short list for clinical examination. The workflow then cleans up the resulting candidate list of identifiable errors and artifacts. Typical candidate lists produced by this pipeline consist of 10-50 variants (Tables S23 and S24). These lists include compound-heterozygous variants located in non-coding regions of the genome.

One new implementation of the workflow is the detection of CNV mosaicism. Mosaicism is a genetic event where a single sample possesses multiple populations of cells that possess different proportions of variants. The goal of the program is to detect stretches of phased variants that show consistent and significant evidence for deviation in allele depth (AD) contributed by the mother and father. The first step is to phase a set of heterozygous genotypes in the proband by examining the parental genotypes. The phasing done here is more stringent than in the previous method described in the vg pedigree workflow because we are looking for a sequence of easily phasable SNPs and so the procedure is rule-based instead of *WhatsHap* which is based on statistical models. A given genotype in the proband is phaseable if three conditions are met: at least one parent has a homozygous genotype, neither parent shares the same genotype, and both alleles are called in both parents genotypes. If a large enough proportion of genotypes are phased in this way, the program examines regions of sufficient length for consecutive stretches of allele balance deviation. A sliding window of 10,000 phased genotypes is used to scan each chromosome and find the boundaries of the mosaic region. For each SNP within this window, the AD of one parent is subtracted from the AD of the other parent. A t-test is applied to the list of AD differences within the window to test if the distribution is significantly different from the null model of no difference. If the t-test statistic is greater than the input threshold, then a region of possible mosaicism is detected and subsequently logged in a separate file for further examination. This threshold was determined empirically against mosaic-positive samples obtained by the UDP. This differs from traditional CNV callers in that this program incorporates trio information to look for partial deletion or duplication events at megabase scales at a continuous level of granularity.

This program can also determine three types of mosaicism: uniparental isodisomy-disomy, trisomy-disomy, and monosomy-disomy. In uniparental isodisomy-disomy mosaicism the individual has populations of cells where a proportion of their genome shares both copies from only one of their parents, and the rest of their cells have inherited a copy from both parents. These types of mosaics are detected by examining the total read depth of the child and parents within the candidate mosaic region. If the proportion of total read depth between the child and parents are the same, and the proportion of ADs of the phasable SNPs between the child and parents are not the same, then the program will classify the mosaic region as uniparental isodisomy-disomy.

In trisomy-disomy mosaicism, the individual has populations of cells where a proportion of their genome has inherited both copies from one of their parents and the rest of their cells have inherited a copy from both parents. If the proportion of total read depth in the child is greater than their parents, then the region is classified as trisomy disomy mosaicism. Alternatively, in monosomy-disomy mosaicism, the individual inherits only one copy from only one parent in some of their cells, and the rest of their cells inherit one copy from each parent. In this case, if the total read depth in the child is less than that of their parents, then the region is classified as monosomy disomy mosaicism.

All modules have been implemented in software containers to improve portability and interoperability with other workflow engines (Kane and Matthias 2018; Schulz et al. 2016). The candidate analysis workflow is implemented within the toil-vg software package under the `toil-vg analysis` subcommand. The candidate analysis workflow is also available in `WDL` format in the Dockstore repository at https://dockstore.org/workflows/github.com/cmarkello/bmtb_wdl/bmtb:main.

## Code Availability and Data Access

Both the VG-Pedigree workflow and the candidate analysis workflow are implemented in the software workflow engine TOIL (Vivian et al. 2017) for cloud-based and cluster-compute systems under the software framework *toil-vg*. They are callable using the `toil-vg pedigree` and `toil-vg analysis` subcommands, respectively. *toil-vg* is available on GitHub at https://github.com/vgteam/toil-vg. The workflows are also made available in `WDL` format in the Dockstore (O’Connor et al. 2017) repository at https://dockstore.org/workflows/github.com/vgteam/vg_wdl/vg-pedigree-giraffe-deeptrio:master and https://dockstore.org/workflows/github.com/cmarkello/bmtb_wdl/bmtb:main.

Input data used in the mapping evaluation, variant calling evaluation and runtime evaluation are all publicly available and listed in the scripts posted in the github repository: https://github.com/cmarkello/vg-pedigree-paper. Input data used in the candidate analysis evaluation experiments have been or are being submitted to the database of Genotypes and Phenotypes (dbGaP).

Scripts for reproducing the methods for graph construction can be found in https://github.com/cmarkello/vg-pedigree-paper/tree/main/scripts/graph_construction. Scripts for reproducing the mapping evaluation experiments can be found in https://github.com/cmarkello/vg-pedigree-paper/tree/main/scripts/wgs_mapping_simulation. Scripts for reproducing the real-data mapping and variant calling evaluation can be found in https://github.com/cmarkello/vg-pedigree-paper/tree/main/scripts/wgs_mapping_experiments and https://github.com/cmarkello/vg-pedigree-paper/tree/main/scripts/wgs_calling_experiments, respectively.

## Acknowledgments

We gratefully thank Dr. William Gahl, Dr. David Adams and the members of the NHGRI Undiagnosed Diseases Program for providing the resources, experiment execution assistance and data access that has made this project possible. We would also like to thank Jouni Sirén, Erik Garrison, Xian Chang, Jean Monlong, Adam Novak and the rest of the Variation Graph team at the UCSC Genomics Institute for providing the tools and methods developed from which much of this work is built upon. All pipelines and evaluations used the computational resources of the NIH HPC Biowulf cluster at the National Institutes of Health, Bethesda, MD (https://hpc.nih.gov).

## Author Contributions

CM, CH, AR, TM and BP planned the analysis. Software and containers were developed by CM, CH, AR, and TM. *DeepVariant* and *DeepTrio* training was planned and executed by AC and PC. CM and TM analyzed the results. CM prepared the manuscript with editing assistance and advice from JE and BP.

